# Estimating time to the common ancestor for a beneficial allele

**DOI:** 10.1101/071241

**Authors:** Joel Smith, Graham Coop, Matthew Stephens, John Novembre

## Abstract

The haplotypes of a beneficial allele carry information about its history that can shed light on its age and putative cause for its increase in frequency. Specifically, the signature of an allele’s age is contained in the pattern of local ancestry that mutation and recombination impose on its haplotypic background. We provide a method to exploit this pattern and infer the time to the common ancestor of a positively selected allele following a rapid increase in frequency. We do so using a hidden Markov model which leverages the length distribution of the shared ancestral haplotype, the accumulation of derived mutations on the ancestral background, and the surrounding background haplotype diversity. Using simulations, we demonstrate how the inclusion of information from both mutation and recombination events increases accuracy relative to approaches that only consider a single type of event. We also show the behavior of the estimator in cases where data do not conform to model assumptions, and provide some diagnostics for assessing and improving inference. Using the method, we analyze population-specific patterns in the 1000 Genomes Project data to provide a global perspective on the timing of adaptation for several variants which show evidence of recent selection and functional relevance to diet, skin pigmentation, and morphology in humans.

## Introduction

A complete understanding adaptation depends on a description of the genetic mechanisms and selective history that underlies adaptive genetic variation [Radwan and Babik, 2012]. Once a genetic variant underlying a putatively adaptive trait has been identified, several questions remain: What is the molecular mechanism by which the variant affects organismal traits and fitness [Dalziel et al., 2009]?; what is the selective mechanism responsible for allelic differences in fitness?; Did the variant arise by mutation more than once [Elmer and Meyer, 2011]?; when did each unique instance of the variant arise and spread [Slatkin and Rannala, 2000]? Addressing these questions for numerous case studies of beneficial variants across multiple species will be necessary to gain full insight into general properties of adaptation. [Stinchcombe and Hoekstra, 2008]

Here, our focus is on the the last of the questions given above; that is, when did a mutation arise and spread? Understanding these dates can give indirect evidence regarding the selective pressure that may underlie the adaptation; This is especially useful in cases where it is logistically infeasible to assess fitness consequences of a variant in the field directly [Barrett and Hoekstra, 2011]. In humans, for example, dispersal across the globe has resulted in the occupation of a wide variety of habitats, and in several cases, selection in response to specific ecological pressures appears to have taken place. There are well-documented cases of loci showing evidence of recent selection in addition to being functionally relevant to known phenotypes of interest [Jeong and Di Rienzo, 2014]. Nakagome et al. (2015) specify time intervals defined by the human dispersal out-of-Africa and the spread of agriculture to show the relative concordance among allele ages for several loci associated with autoimmune protection and risk, skin pigmentation, hair and eye color, and lactase persistence.

When a putative variant is identified as the selected site, the non-random association of surrounding variants on a chromosome can be used to understand its history. This combination of surrounding variants is called a haplotype, and the non-random association between any pair of variants is called linkage disequilibrium (LD). Due to recombination, LD between the focal mutation and its initial background of surrounding variants follows a per-generation rate of decay. New mutations also occur on this haplotype at an average rate per generation. The focal mutation’s frequency follows a trajectory determined by the stochastic outcome of survival, mating success and offspring number. If the allele’s selective benefit increases its frequency at a rate faster than the rate at which LD decays, the resulting signature is one of high LD and a reduction of polymorphism near the selected mutation [Smith and Haigh, 1974, Kaplan et al., 1989]. Many methods to exploit this pattern have been developed in an effort to identify loci under recent positive selection [Tajima, 1989, Fu and Li, 1993, Hudson et al., 1994, Kelly, 1997,Depaulis et al., 1998,Andolfatto et al., 1999, Fay and Wu, 2000, Sabeti et al., 2002, Kim and Stephan, 2002, Kim and Nielsen, 2004, Nielsen et al., 2005, Toomajian et al., 2006, Voight et al., 2006, Tang et al., 2007, Sabeti et al., 2007, Williamson et al., 2007, Pickrell et al., 2009, Chen et al., 2010, Grossman et al., 2013, Chen et al., 2015]. A parallel effort has focused on quantifying specific properties of the signature to infer the age of the selected allele [Serre et al., 1990, Kaplan et al., 1994, Risch et al., 1995, Goldstein et al., 1999, Guo and Xiong, 1997, Slatkin and Rannala, 1997, Stephens et al., 1998, Reich and Goldstein, 1999, Thomson et al., 2000, Slatkin, 2002, Tang et al., 2002, Innan and Nordborg, 2003, Przeworski, 2003, Toomajian et al., 2003, Meligkotsidou and Fearnhead, 2005, Tishkoff et al., 2007, Bryk et al., 2008, Coop et al., 2008, Slatkin, 2008, Peter et al., 2012, Beleza et al., 2013b, Chen and Slatkin, 2013, Chen et al., 2015, Nakagome et al., 2015].

One category of methods used to estimate allele age relies on a point estimate of the mean length of the selected haplotype, or a count of derived mutations within an arbitrary cutoff distance from the selected site [Thomson et al., 2000, Tang et al., 2002, Meligkotsidou and Fearnhead, 2005, Hudson, 2007, Coop et al., 2008]. These approaches ignore uncertainty in the extent of the selected haplotype on each chromosome, leading to inflated confidence in the point estimates.

An alternative approach that is robust to uncertainty in the selected haplotype employs an Approximate Bayesian Computation (ABC) framework to identify the distribution of ages that are consistent with the observed data [Tavaré et al., 1997, Pritchard et al., 1999, Beaumont et al., 2002, Przeworski, 2003, Voight et al., 2006,Tishko_ et al., 2007, Beleza et al., 2013b, Peter et al., 2012, Nakagome et al., 2015]. Rather than model the haplotype lengths explicitly, these approaches aim to capture relevant features of the data through the use of summary statistics. These approaches provide a measure of uncertainy induced by the randomness of recombination, mutation, and genealogical history and produce an approximate posterior distribution on allele age. Despite these advantages, ABC approaches suffer from an inability to capture all relevant features of the sample due to their reliance on summary statistics.

As full-sequencing data become more readily available, defining the summary statistics which capture the complex LD among sites and the subtle differences between haplotypes will be increasingly challenging. For this reason, efficiently computable likelihood functions that leverage the full information in sequence data are increasingly favorable.

Several approaches attempt to compute the full likelihood of the data using an importance sampling framework [Slatkin, 2001, Coop and Griffiths, 2004, Slatkin, 2008, Chen and Slatkin, 2013]. Conditioning on the current frequency of the selected allele, frequency trajectories and genealogies are simulated and given weight proportional to the probability of their occurrence under a population genetic model. While these approaches aim to account for uncertainty in the allele’s frequency trajectory and genealogy, they remain computationally infeasible for large samples or do not consider recombination across numerous loci.

In a related problem, early likelihood-based methods for disease mapping have modelled recombination around the ancestral haplotype, providing information for the time to the common ancestor (TMRCA) rather than time of mutation [Rannala and Reeve, 2001, Rannala and Reeve, 2003, McPeek and Strahs, 1999, Morris et al., 2000, Morris et al., 2002]. These models allowed for the treatment of unknown genealogies and background haplotype diversity before access to large data sets made computation at the genome-wide scale too costly. Inference is performed under Markov chain Monte Carlo (MCMC) to sample over the unknown genealogy while ignoring LD on the background haplotypes, or approximating it using a first-order Markov chain. In a similar spirit, Chen and Slatkin (2015) revisit this class of models to estimate the strength of selection and time of mutation for an allele under positive selection using a hidden Markov model.

Hidden Markov models have become a routine tool for inference in population genetics. The Markov assumption allows for fast computation and has proven an effective approximation for inferring the population-scaled recombination rate, the demographic history of population size changes, and the timing and magnitude of admixture events among genetically distinct populations [Li and Stephens, 2003, Li and Durbin, 2011, Price et al., 2009, Hinch et al., 2011, Wegmann et al., 2011]. The approach taken by Chen and Slatkin (2015) is a special case of two hidden states—the ancestral and background haplotypes. The ancestral haplotype represents the linked background that the focal allele arose on, while the background haplotypes represent some combination of alleles that recombine with the ancestral haplotype during its increase in frequency. Chen and Slatkin (2015) compute maximum-likelihood estimates for the length of the ancestral haplotype on each chromosome carrying the selected allele. Inference for the time of mutation is performed on these fixed estimates assuming they are known. The authors condition the probability of an ancestry switch event on a logistic frequency trajectory for the selected allele and assume independence among haplotypes leading to the common ancestor. The likelihood for background haplotypes is approximated using a first-order Markov chain to account for non-independence among linked sites.

Our approach differs in several ways from that of Chen and Slatkin (2015). First, our method implements an MCMC which samples over the unknown ancestral haplotype to generate a sample of the posterior distribution for the TMRCA instead of the time since mutation. Rather than directly estimate the recombination breakpoints, we integrate over uncertainty among all possible recombination events for each haplotype in the sample. Our model does not assume any particular frequency trajectory, but makes the simplification that all recombination events result in a switch from the ancestral haplotype to the background haplotypes. To incorporate information from derived mutations as well as LD decay, we model differences from the ancestral haplotype as mutation events having occurred since the common ancestor. Rather than use a first-order Markov chain, our emission probabilities account for the LD structure among background haplotypes using the Li and Stephens (2003) haplotype copying model and a reference panel of haplotypes without the selected allele [Li and Stephens, 2003] (Figure 1b,c). The copying model provides an approximation to the coalescent with recombination by modelling the sequence of variants following the recombination event as an imperfect mosaic of haplotypes in the reference panel. Below, we use simulation to show the sensitivity of our model to these simplified assumptions for varying strengths of selection, final allele frequencies, and sampling regimes for the choice of reference panel. An R package is available to implement this method on github (https://github.com/joelhsmth/startmrca).

**Fig. 1.**
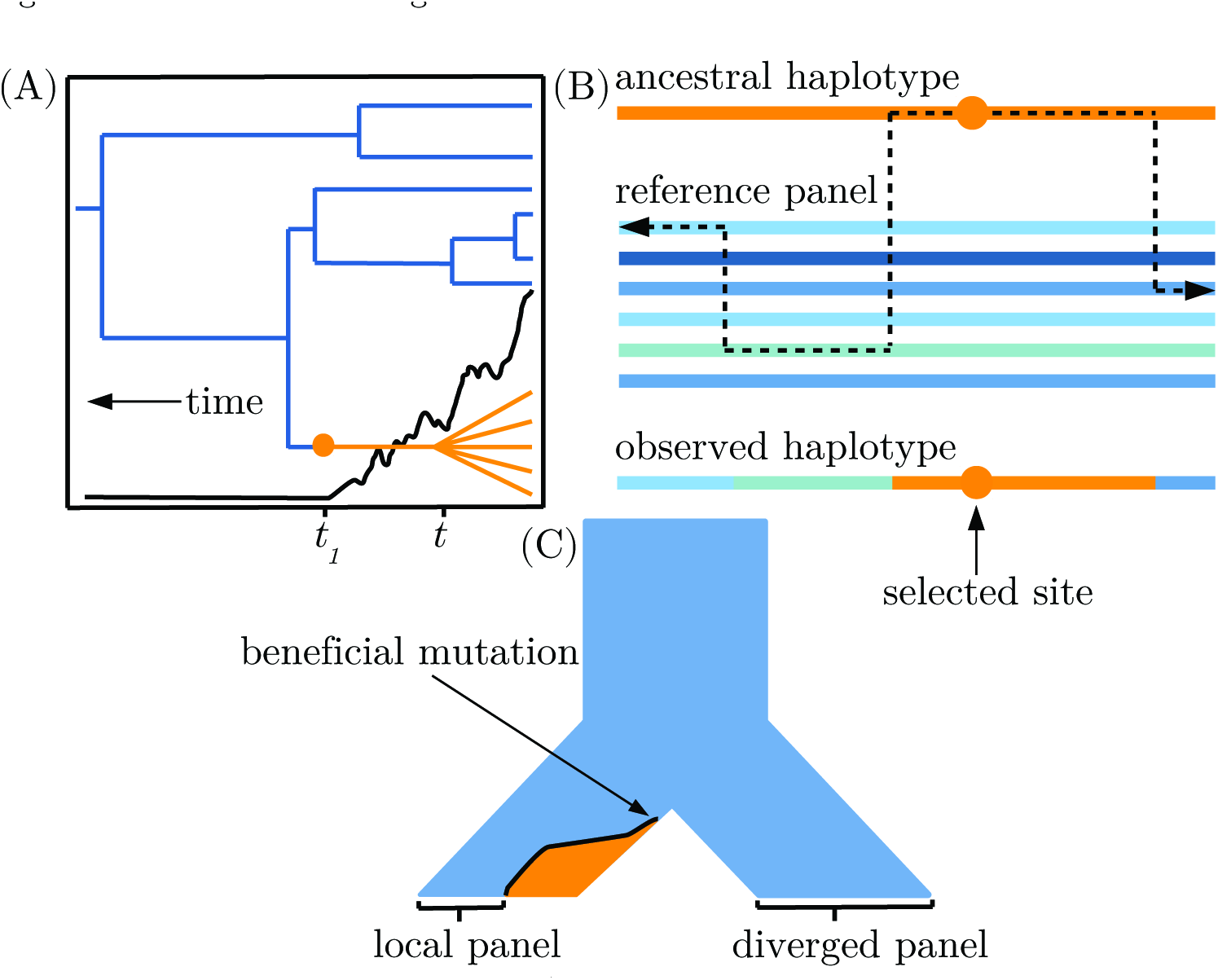
Visual descriptions of the model. a) An idealized illustration of the effect of a selectively favored mutation’s frequency trajectory (black line) on the shape of a genealogy at the selected locus. The orange lineages are chromosomes with the selected allele. The blue lineages indicate chromosomes that do not have the selected allele. Note the discrepancy between the time to the common ancester of chromosomes with the selected allele, *t*, and the time at which the mutation arose, *t*_1_. b) The copying model follows the ancestral haplotype (orange) moving away from the selected site until recombination events within the reference panel lead to a mosaic of non-selected haplotypes surrounding the ancestral haplotype. c) A demographic history with two choices for the reference panel: local and diverged. After the ancestral population at the top of the figure splits into two sister populations, a beneficial mutation arises and begins increasing in frequency. The orange and blue colors indicate frequency of the selected and non-selected alleles, respectively.

## Materials and Methods

### Model description

In general, the TMRCA for a sample of haplotypes carrying the advantageous allele (hereafter referred to as *t*) will be more recent than the time of mutation [Kaplan et al., 1989]. We aim to estimate *t* in the case where a selectively advantageous mutation occured in an ancestor of our sample *t*_1_ generations ago (Fig 1a). Viewed backwards in time, the selected variant decreases in frequency at a rate proportional to the selection strength. For a rapid drop in allele frequency, the coalescent rate among haplotypes carrying the selected variant is amplified. The same effect would be observed for population growth from a small initial size forward in time [Hudson et al., 1990, Slatkin and Hudson, 1991]. As a result, the genealogy of a sample having undergone selection and/or population growth becomes more “star-shaped” (Fig 1a). This offers some convenience, as it becomes more appropriate to invoke an assumption of independence among lineages when selection is strong.

We assume no crossover interference between recombination events within a haplotype, and therefore treat each side flanking the focal allele separately. We define one side of the selected site, within a window of some predetermined length, to have *L* segregating sites, such that an individual’s sequence will be indexed from site *s* = {1,…, *L*}, where s=1 refers to the selected site (a notation reference is provided in Table 1). To simplify notation, this description will be written for a window on one side flanking the selected site. Note that the opposing side of the selected site is modelled in an identical fashion after redefining *L*.

**Table 1.** Notation used to describe the model

|  |  |
| --- | --- |
| $n$ | Number of haplotypes with the selected allele |
| $m$ | Number of haplotypes without the selected allele |
| $L$ | Number of SNPs flanking the selected site (one side considered at a time) |
| $X$ | $n \times L$ matrix of haplotypes with the selected allele |
| $H$ | $m \times L$ matrix of haplotypes without the selected allele |
| $X_{ij}$ | Allele in haplotype $i$ at SNP $j$, where $i \in \{1, \dots, n\}$, and $j \in \{1, \dots, L\}$ |
| $H_{zj}$ | Allele in haplotype $z$ at SNP $j$, where $z \in \{1, \dots, m\}$, and $j \in \{1, \dots, L\}$ |
| $A_j$ | Ancestral allele at site $j$ |
| $Z_j$ | The reference panel haplotype from which $X_i$ copies at site $j$ |
| $t$ | Time to the most recent common ancestor (TMRCA) |
| $W_i$ | The location of the first recombination event off of the ancestral haplotype |
| $r$ | Recombination rate per basepair per generation |
| $\mu$ | Mutation rate per basepair per generation |
| $\theta$ | Haplotype miscopying rate, or population-scaled mutation rate ($4N\mu$) |
| $\rho$ | Haplotype switching rate, or population-scaled recombination rate ($4Nr$) |
| $d_w$ | Physical distance of site $w$ from the selected site, where $w \in \{1, \dots, L\}$ |
| $c_j$ | Number of basepairs between sites $j$ and $j + 1$ |
| $\alpha_{iw}$ | Likelihood of haplotype $i$ for sites $1, \dots, w$ |
| $\beta_{iw}$ | Likelihood of haplotype $i$ for sites $(w + 1), \dots, L$ |

Let *X* denote an *n* × *L* data matrix for a sample of *n* chromosomes with the selected variant. *X_ij_* is the observed allelic type in chromosome *i* at variant site *j*, and is assumed to be biallelic where *X_ij_* ∈ {1, 0}. Let *H* denote an *m* × *L* matrix comprising *m* chromosomes that do not have the selected variant where *H_ij_* ∈ {1, 0}. Let *A* denote the ancestral haplotype as a vector of length *L* where *A_j_* is the allelic type on the ancestral selected haplotype at segregating site *j* and *A_j_* ∈ {1, 0}. We assume independence among lineages leading to the most recent common ancestor of the selected haplotype. This is equivalent to assuming a star-shaped genealogy which, as noted above, is a reasonable assumption for sites linked to a favorable variant under strong selection. We can then write the likelihood as

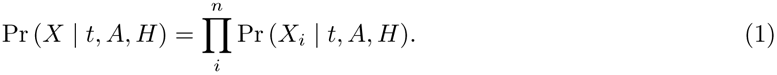

In each individual haplotype, *X_i_*, we assume the ancestral haplotype extends from the selected allele until a recombination event switches ancestry to a different genetic background. Let *W* ∈ {1,…, *L*} indicate that location of the first recombination event occurs between sites *W* and *W* + 1 (*W* = *L* indicates no recombination up to site *L*). We can then condition the probability of the data on the interval where the first recombination event occurs and sum over all possible intervals to express the likelihood as

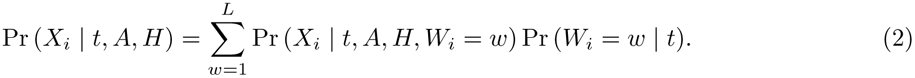

Assuming haplotype lengths are independent and identically distributed draws from an exponential distribution, the transition probabilities for a recombination event off of the ancestral haplotype are

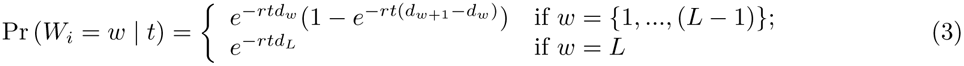

where *d_w_* is the distance, in base pairs, of site *w* from the selected site and *r* is the local recombination rate per base pair, per generation. The data for each individual, *X_i_*, can be divided into two parts: one indicating the portion of an individual’s sequence residing on the ancestral haplotype (before recombining between sites *w* and *w* + 1), *X*_*i*(*j*≤*w*)_, and that portion residing off of the ancestral haplotype after a recombination event, *X*_*i*(*j*>*w*)_. We denote a seperate likelihood for each portion

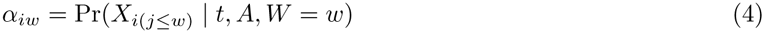

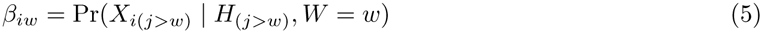

Because the focal allele is on the selected haplotype, *α*_1_ = 1. Conversely, we assume a recombination event occurs at some point beyond locus *L* such that *β_L_* = 1. We model *α* by assuming the waiting time to mutation at each site on the ancestral haplotype is exponentially distributed with no reversal mutations and express the likelihood as

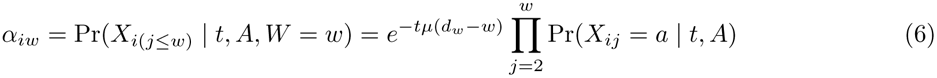

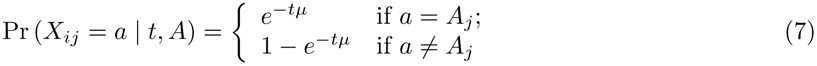

The term, *e*^−*tµ*(*dw*−*w*)^, on the right side of Eq (6) captures the lack of mutation at invariant sites between each segregating site. Assuming *tμ* is small, Eq (6) is equivalent to assuming a Poisson number of mutations (with mean *tμ*) occurring on the ancestral haplotype.

For *β_w_*, the probability of observing a particular sequence after recombining off of the ancestral haplotype is dependent on standing variation in background haplotype diversity. The Li and Stephens (2003) haplotype copying model allows for fast computation of an approximation to the probability of observing a sample of chromosomes related by a genealogy with recombination. Given a sample of *m* haplotypes, *H* ∈ {*h*_1_,…, *h_m_*}, a population scaled recombination rate *ρ* and mutation rate *θ*, an observed sequence of alleles is modelled as an imperfect copy of any one haplotype in the reference panel at each SNP. Let *Z_j_* denote which haplotype *X_i_* copies at site *j*, and *c_j_* denote the number of base pairs between sites *i* and *j*. *Z_j_* follows a Markov process with transition probabilities

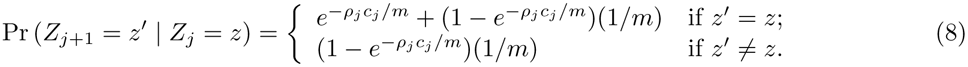

To include mutation, the probability that the sampled haplotype matches a haplotype in the reference panel is *m*/(*m* + *θ*), and the probability of a mismatch (or mutation event) is *θ*/(*m* + *θ*). Letting a refer to an allele where a *a* ∈{1,0}, the matching and mismatching probabilities are

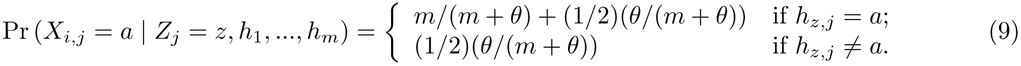

Eq (5) requires a sum over the probabilities of all possible values of *Z_j_* using Eq (8) and Eq (9). This is computed using the forward algorithm as described in Rabiner (1989) and Appendix A of Li and Stephens (2003) [Rabiner, 1989, Li and Stephens, 2003].

The complete likelihood for our problem can then be expressed as

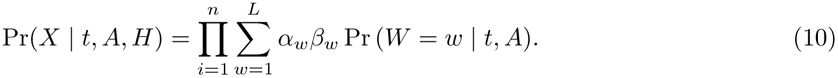

This computation is on the order 2*Lnm*^2^, and in practice for *m* = 20, *n* = 100 and *L* = 4000 takes approximately 3.027 seconds to compute on an Intel® Core^TM^ i7-4750HQ CPU at 2.00GHz × 8 with 15.6 GiB RAM.

### Inference

Performing inference on *t* requires addressing the latent variables *w* and *A* in the model. Marginalizing over possible values of *w* is a natural summation per haplotype that is linear in *L* as shown above. For *A*, 200 the number of possible values is large (2*^L^*), and so we employ a Metropolis–Hastings algorithm to jointly sample the posterior of *A* and *t*, and then we take marginal samples of t for inference. For the priors, we assign a uniform prior density for *A* such that Pr(*A*) = 1/2*^L^*, and we use an improper prior on *t*, a uniform density on all values greater than a positive constant value *u*. Proposed MCMC updates of the ancestral haplotype, *A*′, are generated by randomly selecting a site in *A* and flipping to the alternative allele. For *t*, proposed values are generated by adding a normally distributed random variable centered at 0: *t*′ = *t* + *N*(0, *σ*^2^). To start the Metropolis-Hastings algorithm, an initial value of *t* is uniformly drawn from a user-specified range of values (10 to 2000 in the applications here). To initialize the ancestral haplotype to a reasonable value, we use a heuristic algorithm which exploits the characteristic decrease in variation near a selected site (see S1 Appendix).

For each haplotype in the sample of beneficial allele carriers, the Li and Stephens (2003) model uses a haplotype miscopying rate *θ*, and switching rate *ρ*, to compute a likelihood term for loci following the recombination event off of the ancestral haplotype. For our analyses, we set *ρ* = 4.4 × 10^−4^ using our simulated values of *r* = 1.1 × 10^−8^ per bp per generation and *N* = 10; 000, where *ρ* = 4*Nr*. Following Li and Stephens (2003) we 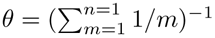; as derived from the expected number of mutation events on a genealogy relating *n* chromosomes at a particular site. We found no discernible effects on estimate accuracy when specifying different values of *ρ* and *θ* (S1 Fig 3, 4).

## Results

### Evaluating accuracy

We generated data using the software mssel (Dick Hudson, personal communication), which simulates a sample of haplotypes conditioned on the frequency trajectory of a selected variant under the structured coalescent [Kaplan et al., 1988, Hudson and Kaplan, 1988]. Trajectories were first simulated forwards in time under a Wright-Fisher model for an additive locus with varying strengths of selection and different ending frequencies of the selected variant. Trajectories were then truncated to end at the first time the allele reaches a specified frequency.

Because our model requires a sample (or “panel’’) of reference haplotypes without the selected allele, we tested our method for cases in which the reference panel is chosen from the local population in which the selected allele is found, as well as cases where the panel is from a diverged population where the selected haplotype is absent (Fig 1c). Regardless of scenario, the estimates are on average within a factor of 2 of the true value, and often much closer. When using a local reference panel, point estimates of *t* increasingly underestimate the true value (TMRCA) as selection becomes weaker and the allele frequency increases (Fig 2). Put differently, the age of older TMRCAs tend to be underestimated with local reference panels. Using the mean posteriors as point estimates, mean values of log_2_(estimate/true value) range from –0.62 to –0.14. Simulations using a diverged population for the reference panel removed the bias, though only in cases where the divergence time was not large. For a reference panel diverged by 0.5*N* generations, mean log_2_(estimate/true value) values range from –0.21 to –0.18. As the reference panel becomes too far diverged from the selected population, estimates become older than the true value (0.36 to 0.94). In these cases, the HMM is unlikely to infer a close match between background haplotypes in the sample and the reference panel, leading to many more mismatches being inferred as mutation events on the ancestral haplotype and an older estimate of *t*.

**Fig. 2.**
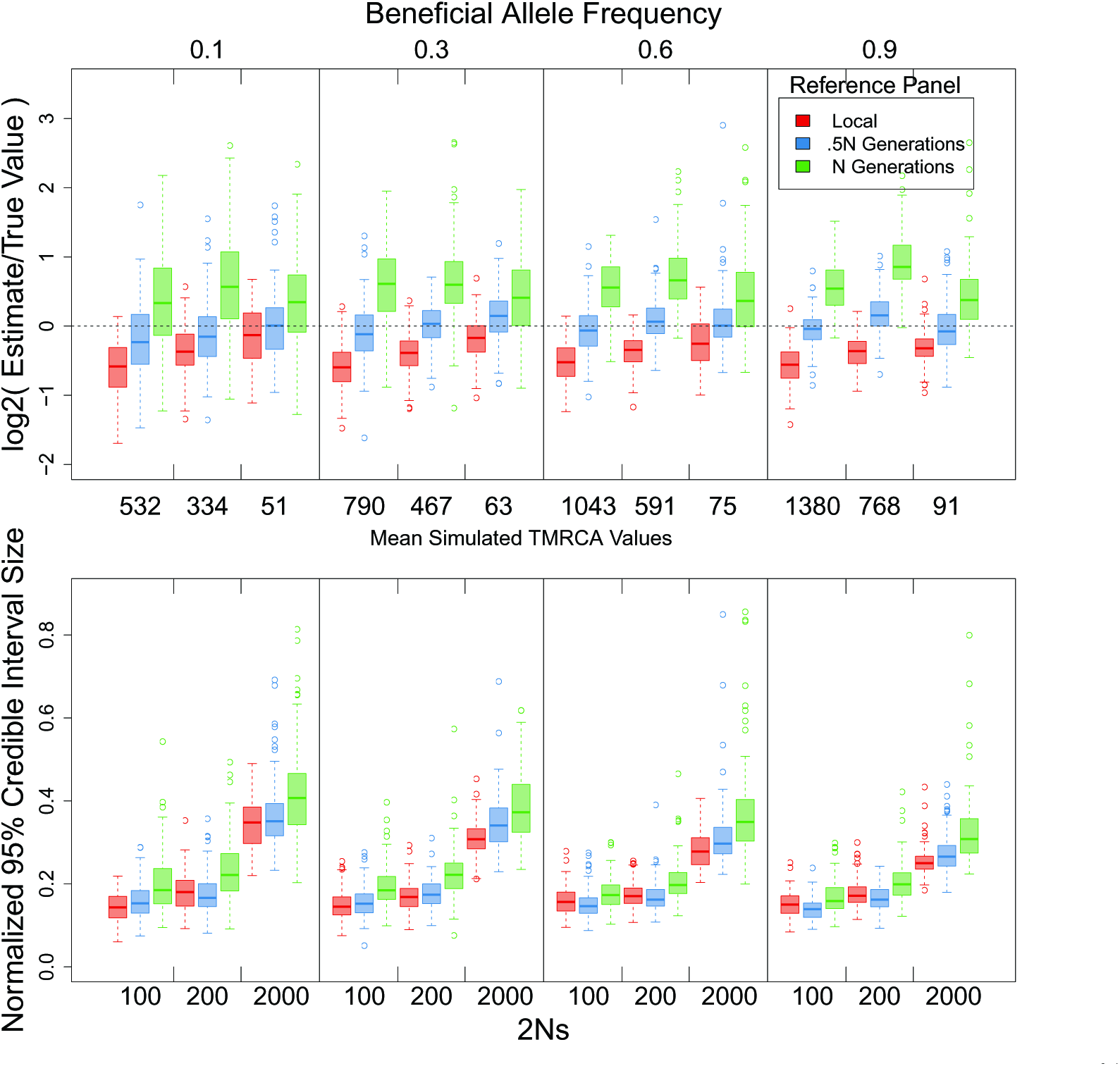
Accuracy results from simulated data. Accuracy of TMRCA point estimates and 95% credible interval ranges from posteriors inferred from simulated data under different strengths of selection, final allele frequencies and choice of reference panel. Credible interval range sizes are in units of generations. 100 simulations were performed for each parameter combination. MCMCs were run for 10000 iterations with a burn-in excluding the first 4000 iterations. A standard deviation of 10 was used for the proposal distribution of *t*. The red boxplots indicate local reference panels. The blue and green boxplots indicate reference panels diverged by .5Ne generations and 1Ne generations, respectively. Each data set was simulated for a 1 Mbp locus with a mutation rate of 1.6 × 10^−8^, recombination rate of 1.1 × 10^−8^ and population size of 10000. Sample sizes for the selected and reference panels were 100 and 20, respectively.

**Fig. 3.**
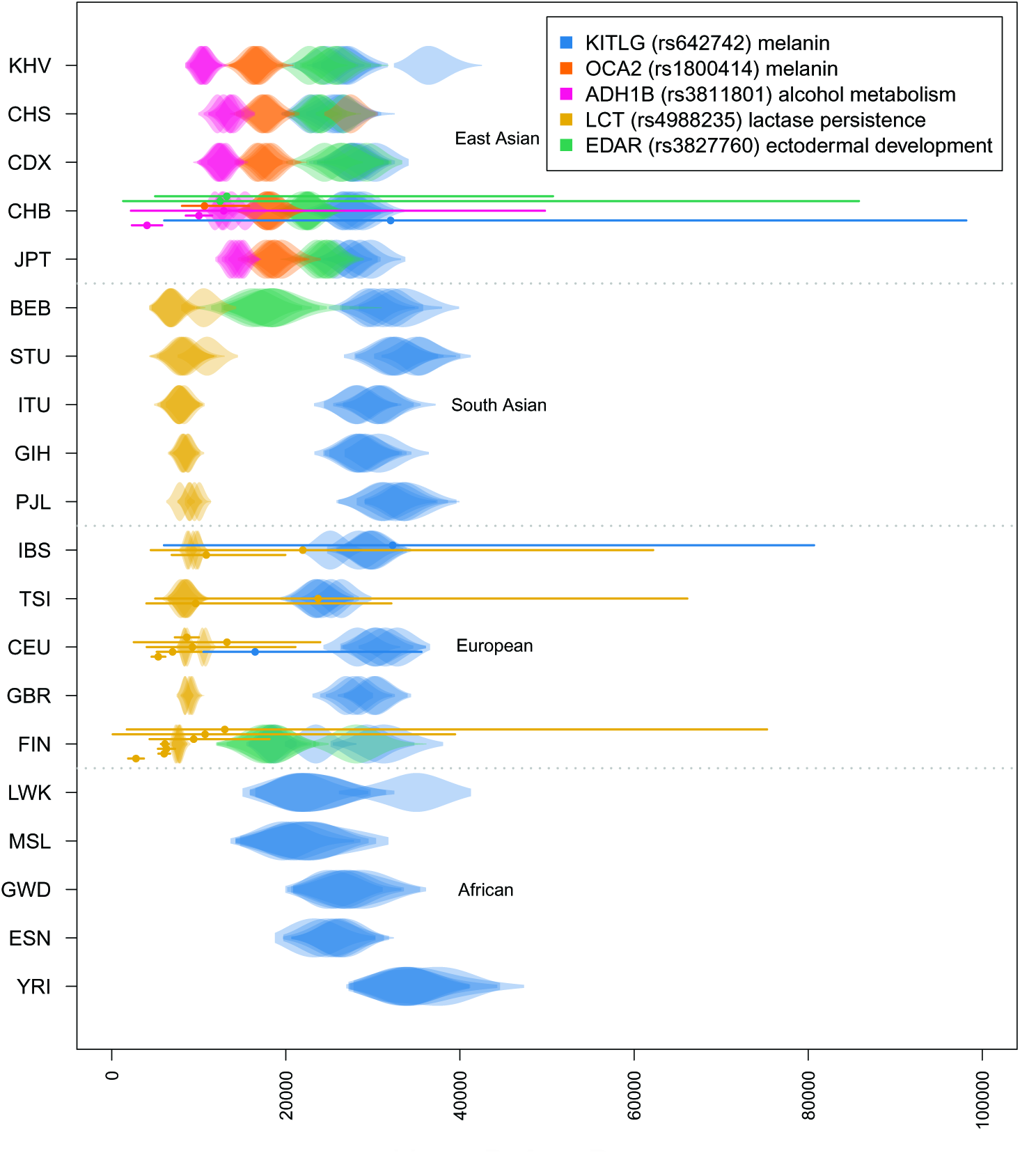
Comparison of TMRCA estimates with previous results. Violin plots of posterior distributions for the complete set of estimated TMRCA values for the 5 variants indicated in the legend scaled to a generation time of 29 years. Each row indicates a population sample from the 1000 Genomes Project panel. Five replicate MCMCs were performed for each variant and population by resampling the selected and reference panels with replacement. Each MCMC was run for 15000 iterations with a standard deviation of 20 for the t proposal distribution. We used the Mb-scale Decode sex-averaged recombination map and a mutation rate of 1.6 × 10^−8^ per basepair per generation. Replicate MCMCs are plotted with transparency. Points and lines overlaying the violins are previous point estimates and 95% confidence intervals for each of the variants indicated by a color and rs number in the legend (see Supplementary Table 1,2). Populations are ordered by broadly defined continental regions. The population sample abbreviations are defined in text.

The bottom panel of Fig 2 shows the effect of selection strength and allele frequency on the size of the normalized 95% credible interval around point estimates. Before normalizing, credible interval sizes using a local reference panel range from 73 to 213 generations for 2*Ns* = 100, versus 18 to 22 generations when 2*Ns* = 2000. Using local and diverged reference panels, we found a minimal effect of the sample size on point estimates (S1 Fig 3, 4). As noted above, higher allele frequencies and weak selection are likely to induce more uncertainty due to the ancestral haplotype tracts recombining within the sample.

To assess the convergence properties of the MCMC, five replicate chains were run for each of 20 simulated data sets produced under three 2*Ns* values (100, 200 and 2000) for frequency trajectories ending at 0.1 (Fig 3).

### Application

We applied our method to five variants previously identified as targets of recent selection in various human populations. Using phased data from the 1000 Genomes Project, we focused on variants that are not completely fixed in any one population so that we could use a local reference panel. The Li and Stephens (2001) haplotype copying model is appropriate in cases where ancestry switches occur among chromosomes within a single population, so we excluded populations in the Americas for which high levels of admixture are known to exist. A brief background on previous work for each locus is provided below. In four of the five cases, variants chosen to be the selected site control observable differences in expression or enzyme activity in functional assays.

In contrast to previous work, we provide age estimates across a range of population samples. This provides greater context to understand the relative timing of adaptation for different populations among each of the selected variants. In addition to the mean TMRCA estimates, the dispersion of estimates for replicate MCMCs using different subsamples of the data can shed light on any genealogical structure in the data and the degree to which a star-genealogy assumption is satisfied. In general, we expect older TMRCAs to have more non-independence represented in the sample’s genealogy and greater dispersion of estimates.

For more efficient run times of the MCMC, we set a maximum number of individuals to include in the selected and reference panels to be 100 and 20, respectively. In cases where the true number of haplotypes for either panel was greater than this in the full data set, we resampled a subset of haplotypes from each population for a total of five replicates per population. For simulation results supporting the use of this resampling strategy, see Supplementary Fig 4. The MCMCs were run for 15000 iterations with 9000 burn-in iterations. Figures 4 and 5 show the results for all five variants along with previous point estimates and 95% confidence intervals assuming a generation time of 29 years [Fenner, 2005]. Supplementary Table 1 lists the mean and 95% credible intervals for estimates with the highest mean posterior probability which we refer to in the text below. Supplementary Table 2 lists the previous estimates and confidence intervals with additional details of the different approaches taken.

**Fig. 4.**
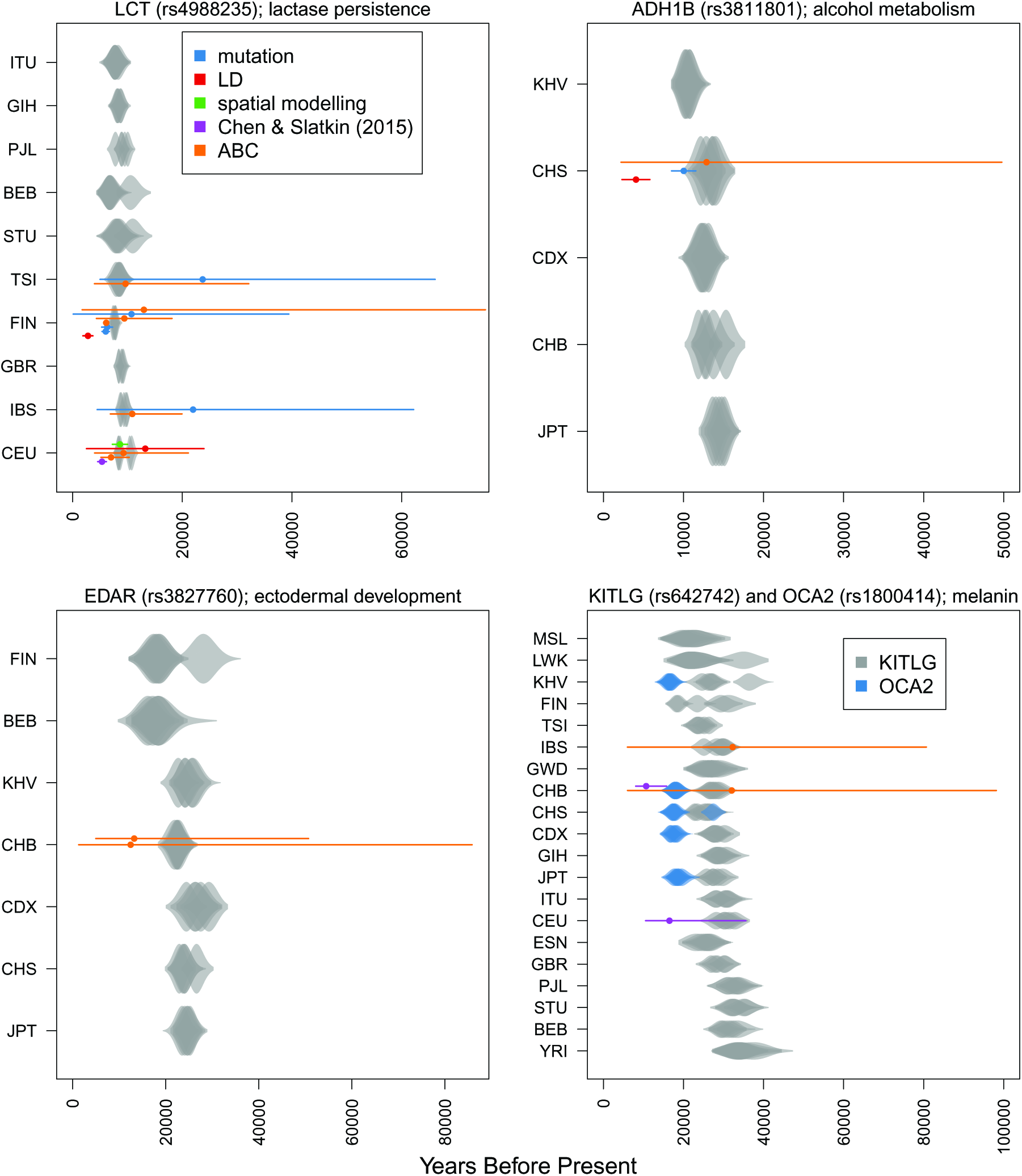
Comparison of TMRCA estimates and previous estimate approaches. Results from Fig 4 sorted into different plots for different variants. Previous estimates are colored by an abbreviated description of the type of information used in the data. The blue violin plots in the KITLG/OCA2 plot are estimates for the OCA2 variant. The purple and orange previous estimates for CHB in the KITLG/OCA2 plot refer to OCA2 and KITLG, respectively.

To model recombination rate variation, we used recombination rates from the Decode sex-averaged recombination map inferred from pedigrees among families in Iceland [Kong et al., 2010]. Because some populations may have recombination maps which differ from the Decode map at fine scales, we used a mean uniform recombination rate inferred from the 1 megabase region surrounding each variant. The motivation for this arises from how recombination rate variation across the genome has been previously shown to remain relatively consistent among recombination maps inferred for different populations at the megabase-scale [Broman et al., 1998, Kong et al., 2002, Kong et al., 2010, Baudat et al., 2010, Auton and McVean, 2012]. Further, we found our estimates depend mostly on having the megabase-scale recombination rate appropriately set, with little difference in most cases for estimates obtained by modeling the full recombination map at each locus (S1 Fig 3). We specify the switching rate among background haplotypes after recombining off of the ancestral 285 haplotype to be 4*Nr*, where *N* = 10; 000 and *r* is the 286 mean recombination rate for the 1Mb locus.

For modeling mutation, a challenge is that previous mutation rate estimates vary depending on the approach used [Scally and Durbin, 2012, Ségurel et al., 2014]. Estimates using the neutral substitution rate between humans and chimps are more than 2 × 10^−8^ per bp per generation, while estimates using whole genome sequencing are closer to 1 × 10^−8^. As a compromise, we specify a mutation rate of 1.6 × 10^−8^.

The population sample abbreviations referred to in all of the figures correspond to the following: CHB (Han Chinese in Bejing, China), JPT (Japanese in Tokyo, Japan), CHS (Southern Han Chinese), CDX (Chinese Dai in Xishuangbanna, China), KHV (Kinh in Ho Chi Minh City, Vietnam), CEU (Utah Residents (CEPH) with Northern and Western European Ancestry), TSI (Toscani in Italia), FIN (Finnish in Finland), GBR (British in England and Scotland), IBS (Iberian Population in Spain) YRI (Yoruba in Ibadan, Nigeria), LWK (Luhya in Webuye, Kenya), GWD (Gambian in Western Divisions in the Gambia), MSL (Mende in Sierra Leone), ESN (Esan in Nigeria), GIH (Gujarati Indian from Houston, Texas), PJL (Punjabi from Lahore, Pakistan), BEB (Bengali from Bangladesh), STU (Sri Lankan Tamil from the UK), ITU (Indian Telugu from the UK).

### ADH1B

A derived allele at high frequency among East Asians at the ADH1B gene (rs3811801) has been shown to be functionally relevant for alcohol metabolism [Osier et al., 2002, Eng et al., 2007]. Previous age estimates are consistent with the timing of rice domestication and fermentation approximately 10,000 years ago [Li et al., 2007, Peng et al., 2010, Peter et al., 2012]. However, a more recent estimate by Peter et al. (2012) pushes this time back several thousand years to 12,876 (2,204 - 49,764) years ago. Our results are consistent with an older timing of selection, as our CHB sample (Han Chinese in Beijing, China) TMRCA estimate is 15,377 (13,763 - 17,281) years. Replicate chains of the MCMC are generally consistent, with the oldest estimates in the CHB sample showing the most variation among resampled datasets and the youngest estimate of 10,841 (9,720 - 12,147) in the KHV sample showing the least (JPT and KHV refer to Japanese in Tokyo, Japan and Kinh in Ho Chi Minh City, Vietnam respectively). When using a Mbp-scale recombination rate, all of the ADH1B TMRCAs are inferred to be slightly younger (S1 Fig 3).

### EDAR

Population genomic studies have repeatedly identified the gene EDAR to be under recent selection in East Asians [Akey et al., 2004, Williamson et al., 2005, Voight et al., 2006] with a particular site (rs3827760) showing strong evidence for being the putative target. Functional assays and allele specific expression differences at this position show phenotypic associations to a variety of phenotypes including hair thickness and dental morphology [Bryk et al., 2008, Fujimoto et al., 2008, Kimura et al., 2009].

Our estimate of 22,192 (19,683 - 25,736) years for the EDAR allele in the CHB sample is older than ABC-based estimates of 12,458 (1,314 - 85,835) and 13,224 (4,899 - 50,692) years made by Bryk et al. (2008) and Peter et al. (2012), respectively. We included all populations for which the variant is present including the FIN and BEB samples where it exists at low frequency. Our results for the youngest TMRCAs are found in these two low frequency populations where the estimate in FIN is 17,386 (13,887 - 324 20,794) and the estimate in BEB is 18,370 (14,325 - 22,872). Among East Asian populations, the oldest and youngest TMRCA estimates are found in the KHV sample (25,683; 23,169 - 28,380) and CHB sample (22,192; 19,683 - 25,736).

### LCT

The strongest known signature of selection in humans is for an allele at the LCT gene (rs4988235) which confers lactase persistence into adulthood–a trait unique among mammals and which is thought to be a result of cattle domestication and the incorporation of milk into the adult diets of several human populations [Hollox et al., 2001, Enattah et al., 2002, Bersaglieri et al., 2004,Tishkoff et al., 2007]. There are multiple alleles that show association with lactase persistence [Tishkoff et al., 2007]. We focused on estimating the age of the T-13910 allele, primarily found at high frequency among Northern Europeans, but which is also found in South Asian populations. In addition to complete association with the lactase persistence phenotype, this allele has been functionally verified by *in vitro* experiments [Kuokkanen et al., 2006, Olds and Sibley, 2003, Troelsen et al., 2003].

Mathieson et al. (2015a) use ancient DNA collected from 83 human samples to get a better understanding of the frequency trajectory for several adaptive alleles spanning a time scale of 8,000 years. For the LCT persistence allele (rs4988235), they find a sharp increase in frequency in the past 4,000 years ago. While this is more recent than previous estimates, an earlier TMRCA or time of mutation is still compatible with this scenario.

Our estimates using European and South Asian samples fall between the range from 5000 to 10,000 years ago, which is broadly consistent with age estimates from modern data. The credible intervals for estimates in all of the samples have substantial overlap which makes any ranking on the basis of point estimates difficult. We infer the PJL (Punjabi from Lahore, Pakistan) sample to have the oldest TMRCA estimate of 9,514 (8,596 - 10,383) years. Itan et al. (2009) use spatial modelling to infer the geographic spread of lactase allele from northern to southern Europe. Consistent with their results, the youngest estimate among European populations is found in the IBS sample at 9,341 (8,688 - 9,989) years. Among all samples, the youngest estimate was found in BEB at 6,869 (5,143 - 8809).

### KITLG and OCA2

The genetic basis and natural history of human skin pigmentation is a well studied system with several alleles of major effect showing signatures consistent with being targets of recent selection [Jablonski and Chaplin, 2012, Beleza et al., 2013b, Wilde et al., 2014, Eaton et al., 2015]. We focused on an allele found at high frequency world-wide among non-African populations at the KITLG locus (rs642742) which shows significant effects on skin pigmentation differences between Europeans and Africans [Miller et al., 2007]; although more recent work fails to find any contribution of KITLG toward variation in skin pigmentation in a Cape Verde African-European admixed population [Beleza et al., 2013a]. We also estimated the TMRCA for a melanin-associated allele at the OCA2 locus (rs1800414) which is only found among East Asian populations at high frequency [Edwards et al., 2010].

For the KITLG variant, our estimates among different populations vary from 18,000 to 34,000 years ago, with the oldest age being in the YRI (Yoruba in Ibadan, Nigeria) sample (33,948; 28,861 - 39,099). The youngest TMRCA is found in FIN at 18,733 years (16,675 - 20,816). The next two youngest estimates are also found in Africa with the TMRCA in the MSL (Mende in Sierra Leone) sample being 22,340 (15,723 - 28,950) years old, and that for LWK (Luhya in Webuye, Kenya) being 22,784 (17,922 - 2,8012) years old, suggesting a more complex history than a model of a simple allele frequency increase outside of Africa due to pigmentation related selection pressures. Previous point estimates using rejection sampling approaches on a Portuguese sample (32,277; 6,003 - 80,683) and East Asian sample (32,045; 6,032 - 368 98,165) are again most consistent with our own results on the IBS (29,731; 26,170 - 32,813) and CHB samples (26,773; 24,297 - 30,141) [Beleza et al., 2013b, Chen et al., 2015]. Among East Asians, the oldest and youngest estimates are again found in the JPT (28,637; 24,297 - 30,141) and KHV (24,544; 21,643 - 371 27,193) samples, respectively. The TMRCA for OCA2 alleles in the JPT (18,599; 16,110 - 20,786) and 372 KHV (16370; 14,439 - 18,102) samples are also the oldest and youngest, respectively. When using a 373 Mbp-scale recombination rate, all of the ADH1B estimates are inferred to be slightly younger (S1 Fig 3) than estimates from the fine-scale map.

## Discussion

Our approach uses a simplified model of haplotype evolution to more accurately estimate the timing of selection on a beneficial allele. By leveraging information from carriers and non-carriers of the allele, we can more effectively account for uncertainty in the extent of the ancestral haplotype and derived mutations. Using simulations, we show the performance of our method for different strengths of selection, beneficial allele frequencies and choices of reference panel. By applying our method to five variants previously identified as targets of natural selection in human populations, we provide a comparison among population-specific TMRCAs. This gives a more detailed account of the order in which populations may have acquired the variant and/or experienced selection for the trait it underlies. Comparisons across variants also help to identify patterns of adaptation that are consistent for particular populations or regions of the world.

In that regard, it is hypothesized that local selection pressures and a cultural shift toward agrarian societies has induced adaptive responses among human populations around the globe. As would be expected for the traits associated with the loci studied here, all of our inferred TMRCA values are more recent than the earliest estimates that are commonly used for the dispersal out-of-Africa 40,000 years ago [Benazzi et al., 2011, Higham et al., 2011, Mellars, 2011]. However, the data associated with some variants seem to indicate more recent selective events than others. Our results for variants associated with dietary traits at the LCT and ADH1B genes both imply relatively recent TMRCAs, consistent with hypotheses that selection on these mutations results from recent changes in human diet following the spread of agriculture [Simoons, 1970, Peng et al., 2010]. In contrast, the inferred TMRCAs for EDAR, KITLG and OCA2 imply older adaptive events which may have coincided more closely with the habitation of new environments or other cultural changes.

Several hypotheses have been suggested to describe the selective drivers of skin pigmentation differences among human populations, including reduced UV radiation at high latitudes and vitamin D deficiency [Loomis, 1967, Jablonski and Chaplin, 2000]. Estimated TMRCAs for the variants at the OCA2 and EDAR loci among East Asians appear to be as young or younger than the KITLG variant, but older than the LCT and ADH1B locus. This suggests a selective history in East Asian populations leading to adaptive responses for these traits occurring after an initial colonization. In some cases, the dispersion of replicate MCMC estimates make it difficult to describe the historical significance of an observed order for TMRCA values. However, the consistency of estimates among different populations for particular variants add some confidence to our model’s ability to reproduce the ages which are relevant to those loci or certain geographic regions.

We also compared our estimates to a compilation of previous age estimates based on the time of mutation, time since fixation, or TMRCA of variants associated with the genes studied here. The range of confidence interval sizes for these studies is largely a reflection of the assumptions invoked or relaxed for any one method, as well as the sample size and quality of the data used. Relative to the ABC approaches which are most commonly used today, our method provides a gain in accuracy while accounting for uncertainty in both the ancestral haplotype and its length on each chromosome. Notably, our method provides narrower credible intervals by incorporating the full information from ancestral haplotype lengths, derived mutations, and a reference panel of non-carrier haplotypes.

One caveat of our method is its dependence on the reference panel, which is intended to serve as a representative sample of non-ancestral haplotypes in the population during the selected allele’s increase in frequency. Four possible challenges can arise: (1) segments of the ancestral selected haplotype may be present in the reference panel due to recombination, (this is more likely for alleles that have reached higher frequency), (2) the reference panel may contain haplotypes that are similar to the ancestral haplotype due to low levels of genetic diversity, (3) the reference panel may be too diverged from the focal population, and (4) population connectivity and turnover may lead the “local” reference panel to be largely composed of migrant haplotypes which were not present during the selected allele’s initial increase in frequency.

Under scenarios 1 and 2, the background haplotypes will be too similar to the ancestral haplotype and it may be difficult for the model to discern a specific ancestry switch location. This leads to fewer differences (mutations) than expected between the ancestral haplotype and each beneficial allele carrier. The simulation results are consistent with this scenario: our method tends to underestimate the true age across a range of selection intensities and allele frequencies when using a local reference panel.

Conversely, under scenarios 3 and 4 the model will fail to describe a recombinant haplotype in the sample of beneficial allele carriers as a mosaic of haplotypes in the reference panel. As a result, the model will infer more mutation events to explain observed differences from the ancestral haplotype. Our simulation results show this to be the case with reference panels diverged by *N* generations: posterior mean estimates are consistently older than their true value. Our simulations are perhaps pessimistic though - we chose reference panel divergence times of *N* and 0.5*N* generations, approximately corresponding to *F_ST_* values of 0.4 and 0.2, respectively. For the smaller *F_ST_* values observed in humans, we expect results for diverged panels to closer to those obtained with the local reference panel. Nonetheless, future extensions to incorporate multiple populations within the reference panel would be helpful and possible by modifying the approach of Price et al. (2009). Such an approach would also enable the analysis of admixed populations (we excluded admixed samples from our analysis of the 1000 Genomes data above).

Aside from the challenges imposed by the choice of reference panel, another potential source of bias lies in our transition probabilities, which are not conditioned on the frequency of the selected variant. In reality, recombination events at some distance away from the selected site will only result in a switch from the ancestral to background haplotypes at a rate proportional to 1–*p_l_*, where *p_l_* is the frequency of the ancestral haplotype alleles at locus *l*. In this way, some recombination events may go unobserved – as the beneficial allele goes to high frequency the probability of an event leading to an observable ancestral to background haplotype transition decreases. One solution may be to include the frequency-dependent transition probabilities derived by Chen and Slatkin (2015). Under their model, the mutation time is estimated by assuming a deterministic, logistic frequency trajectory starting at 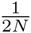. One concern is the specification of a initial frequency for our case, which should correspond to the frequency at which the TMRCA occurs rather than time of mutation. Griffiths and Tavare (1994) derive a framework to model a genealogy under arbitrary population size trajectories, which should be analogous to the problem of an allele frequency trajectory, and other theory on intra-allelic genealogies may be useful here as well [Griffiths and Tavare, 1994, Wiuf and Donnelly, 1999, Wiuf, 2000, Slatkin and Rannala, 2000]. An additional benefit of using frequency trajectories would be the ability to infer posterior distributions on selection coefficients.

Our model also assumes independence among all haplotypes in the sample in a composite-likelihood framework, which is equivalent to assuming a star-genealogy [Varin et al., 2011, Larribe and Fearnhead, 2011]. This is unlikely to be the case when sample sizes are large or the TMRCA is old. It is also unlikely to be true if the beneficial allele existed on multiple haplotypes preceding the onset of selection, was introduced by multiple migrant haplotypes from other populations, or occurred by multiple independent mutation events [Innan and Kim, 2004, Hermisson and Pennings, 2005, Prezeworski et al., 2005, Pritchard et al., 2010, Berg and Coop, 2015].

If the underlying allelic genealogy is not star-like, one can expect different estimates of the TMRCA for different subsets of the data. We suggest performing multiple MCMCs on resampled subsets of the data to informally diagnose whether there are violations from the star-like genealogy assumption. We speculate that exactly how the TMRCAs vary may provide insight to the underlying history. In cases where the TMRCA estimates for a particular population are old and more variable than other populations, the results may be explained by structure in the genealogy, whereby recent coalescent events have occurred among the same ancestral haplotype before the common ancestor. When estimates are dispersed among resampled datasets, in addition to being relatively young, the presence of multiple ancestral haplotypes prior to the variant’s increase in frequency may be a better explanation. Further support for this explanation might come from comparisons to other population samples which show little to know dispersion of estimates from resampled datasets. Future work might make it possible to formalize this inference process.

One possible future direction may be to explicitly incorporate the possibility of multiple ancestral haplotypes within the sample. Under a disease mapping framework, Morris et al. (2002) implement a similar idea in the case where independent disease causing mutations arise at the same locus leading to independent genealogies, for which they coin the term “shattered coalescent’’. For our case, beneficial mutations may also be independently derived on different haplotypes. Alternatively, a single mutation may be old enough to reside on different haplotypes due to a sufficient amount of linked variation existing prior to the onset of selection. Berg and Coop (2015) model selection from standing variation to derive the distribution of haplotypes that the selected allele is present on.

While we have treated the TMRCA as a parameter of interest, our method also produces a sample of the posterior distribution on the ancestral haplotype. This could provide useful information to estimate the frequency spectrum of derived mutations on the ancestral haplotype. Such information could shed light on the genealogy and how well it conforms to the star-shape assumption. The extent of the ancestral haplotype in each individual may also prove useful for identifying deleterious alleles that have increased in frequency as a result of strong positive selection on linked beneficial alleles [Chun and Fay, 2011, Hartfield and Otto, 2011]. For example, Huff et al. (2012) describe a risk allele for Crohn’s disease at high frequency in European populations which they suggest is linked to a beneficial allele under recent selection. Similar to an admixture mapping approach, our method could be used to identify risk loci by testing for an association between the ancestral haplotype and disease status. As another application, identifying the ancestral haplotype may be useful in the context of identifying a source population (or species) for a beneficial allele prior to its introduction and subsequent increase in frequency in the recipient population.

In many cases, the specific site under selection may be unknown or restricted to some set of putative sites. While our method requires the position of the selected site be specified, future extensions could treat the selected site as a random variable to be estimated under the same MCMC framework. This framework would also be amenable to marginalizing over uncertainty on the selected site.

While we focus here on inference from modern DNA data, the increased accessibility of ancient DNA has added a new dimension to population genetic datasets [Lazaridis et al., 2014, Skoglund et al., 2014, Allentoft et al., 2015, Haak et al., 2015, Mathieson et al., 2015a, Mathieson et al., 2015b]. Because it will remain difficult to use ancient DNA approaches in many species with poor archaeological records, we believe methods based on modern DNA will continue to be useful going forward. That said, ancient DNA are providing an interesting avenue for comparative work between inference from modern and ancient samples. For example, Nakagome et al. (2015) use simulations to assess the fit of this ancient DNA polymorphism to data simulated under their inferred parameter values for allele age and selection intensity and they find reasonable agreement. Much work still remains though to fully leverage ancient samples into population genetic inference while accounting for new sources of uncertainty and potential for sampling bias.

Despite these challenges, it is clear that our understanding of adaptive history will continue to benefit from new computational tools which extract insightful information from a diverse set of data sources.

## Acknowledgements

We would like to thank Hussein Al-Asadi, Arjun Biddanda, Anna Di Rienzo, Dick Hudson, Choongwon Jeong, Evan Koch, Joseph Marcus, Shigeki Nakagome, Ben Peter, Mark Reppel, Alex White, members of the Coop lab at UC Davis, members of the Przeworski lab at Columbia University, and members of the He and Stephens labs at the University of Chicago for helpful comments. JS was supported by an NSF Graduate Research Fellowship and National Institute Of General Medical Sciences of the National Institutes of Health under award numbers DGE-1144082 and T32GM007197, respectively. This work was also supported by the National Institute of General Medical Sciences of the National Institutes of Health under award numbers RO1GM83098 and RO1GM107374 to GC, as well as NIH ROI (R01HG007089) to 520 JN.

## Supporting Information

**S1 Fig.**
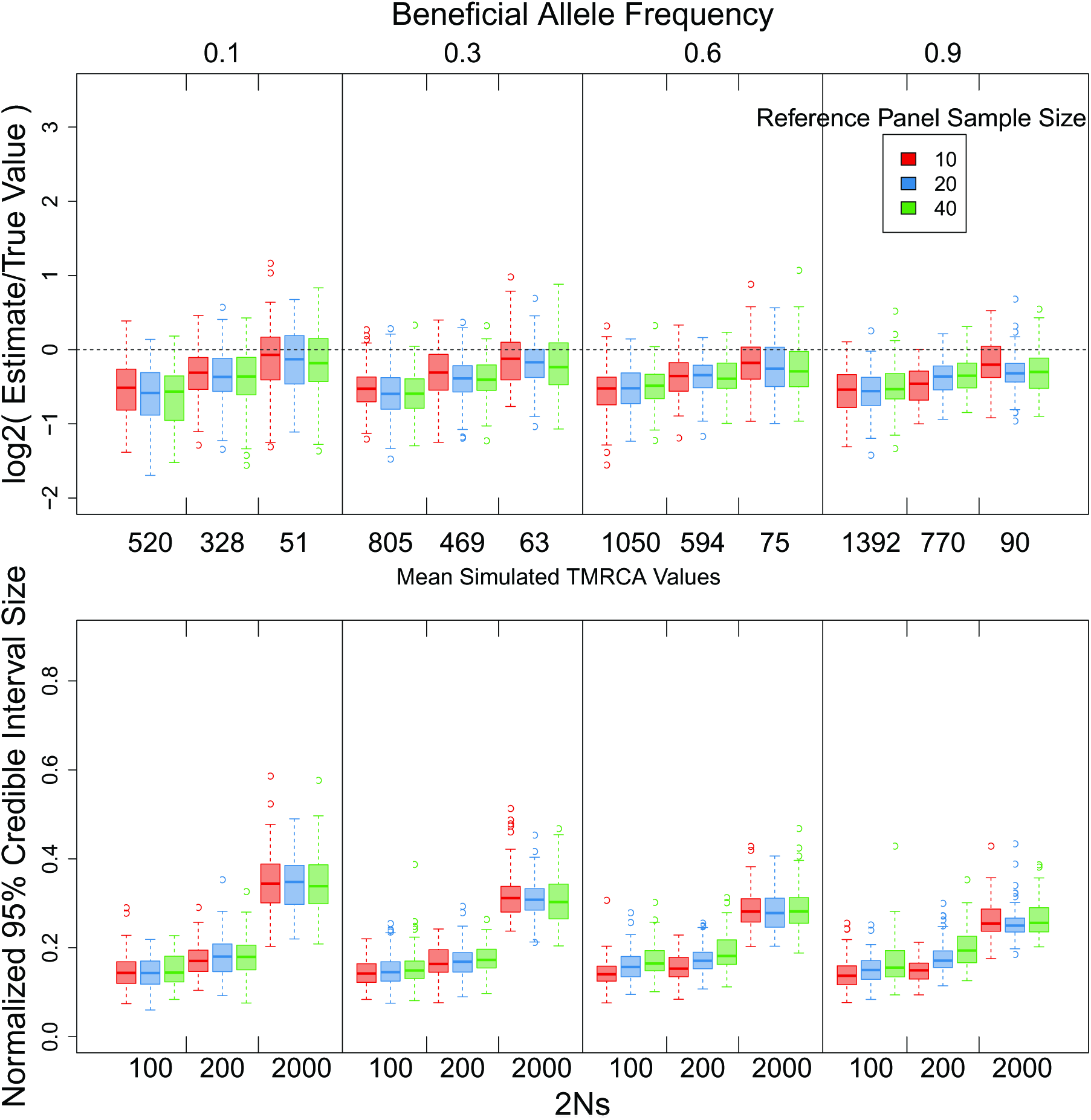
Effect of local reference panel sample size on estimate accuracy. Accuracy of point estimates and 95% credible interval ranges from posteriors inferred from simulated data under different strengths of selection, final allele frequencies and sample sizes for a reference panel. In all cases the reference panel is sampled from the local population where the selected allele is found. All other parameter values are identical to Figure 2 in the main text.

**S2 Fig.**
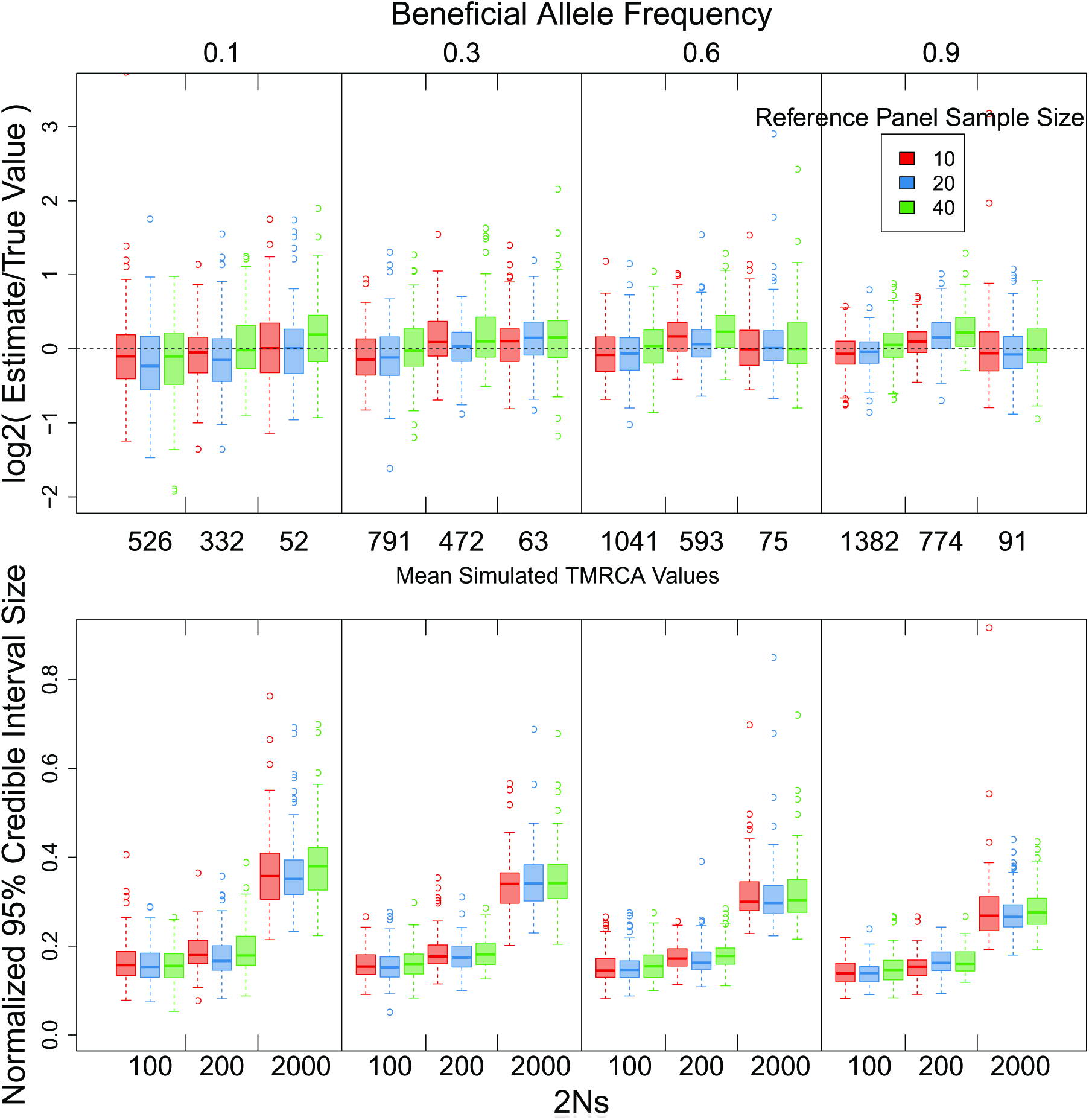
Effect of diverged reference panel sample size on estimate accuracy. Accuracy of point estimates and 95% credible interval ranges from posteriors inferred from simulated data under different strengths of selection, final allele frequencies and sample sizes for a reference panel. In all cases the reference panel is sampled from a population 0.5Ne generations diverged from the selected population. All other parameter values are identical to Figure 2 in the main text.

**S3 Fig.**
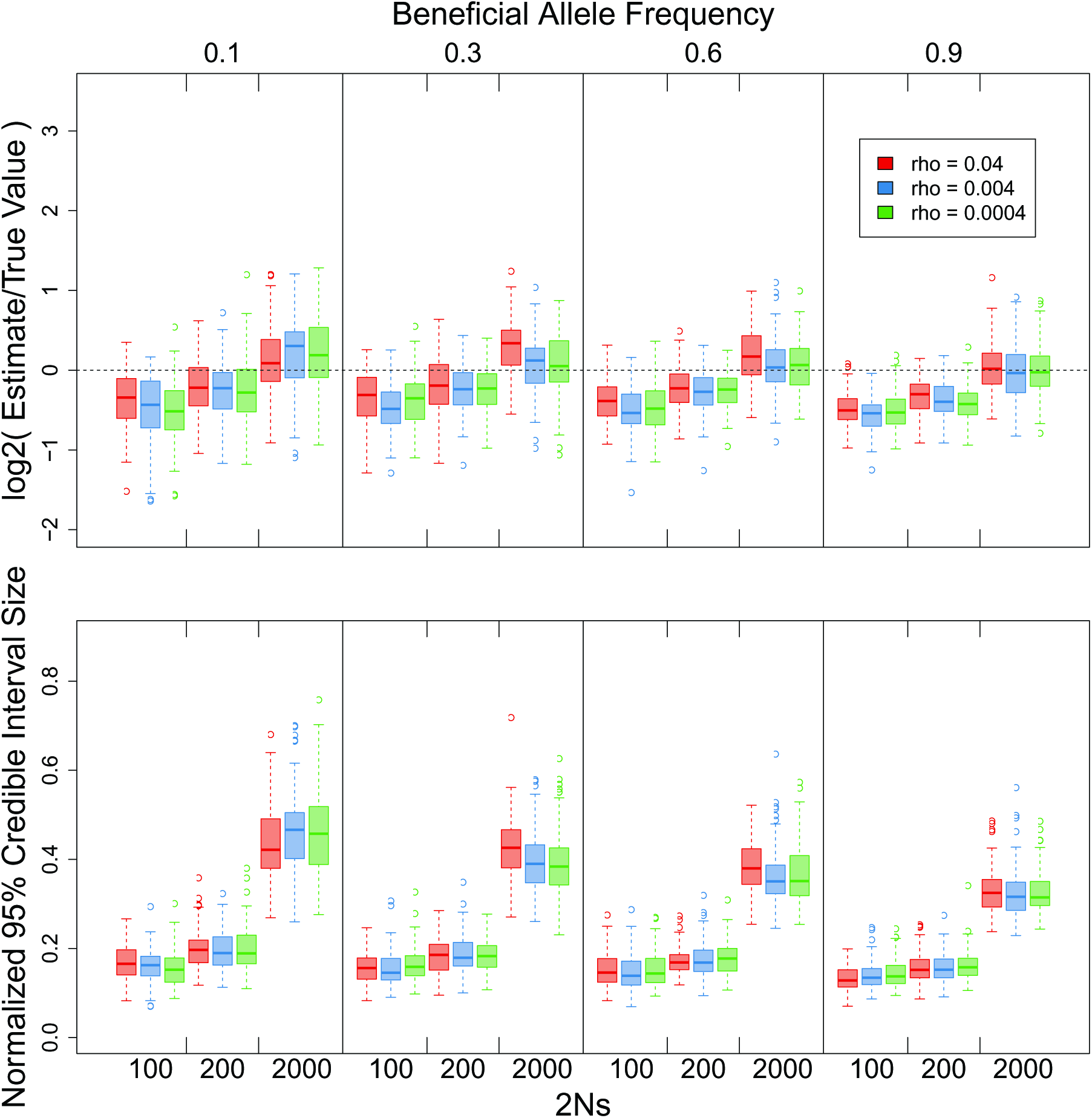
Effect of 535 misspecifying *ρ*. Accuracy results for 3 different values of *ρ* used in the Li and Stephens (2003) copying model for background haplotypes in a local reference panel. All other parameter values are identical to Figure 2 in the main text.

**S4 Fig.**
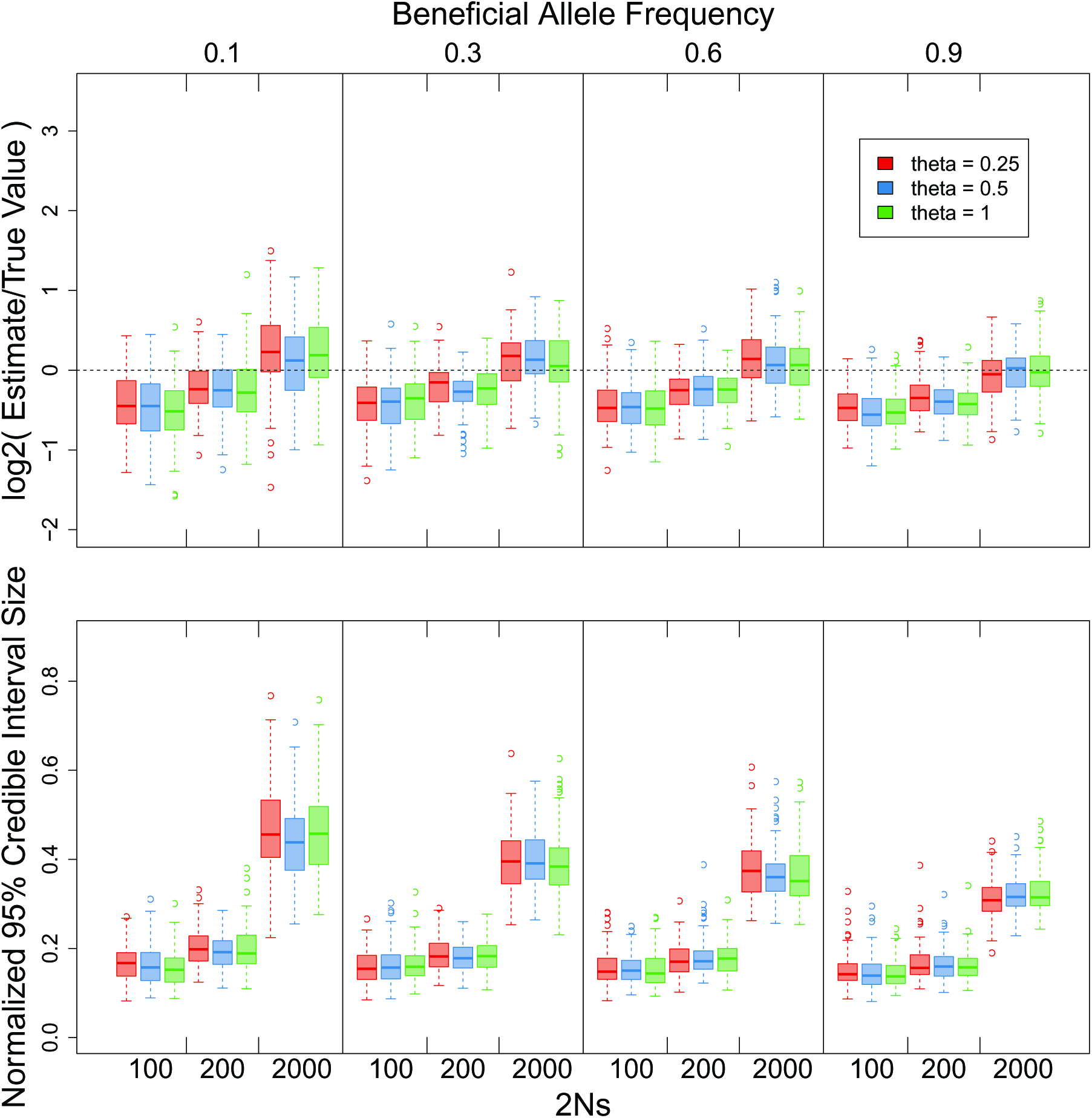
Effect 539 of misspecifying *θ*. Accuracy results for 3 different values of *θ* used in the Li and Stephens (2003) copying model for background haplotypes in a local reference panel. All other parameter values are identical to Figure 2 in the main text.

**S5 Fig.**
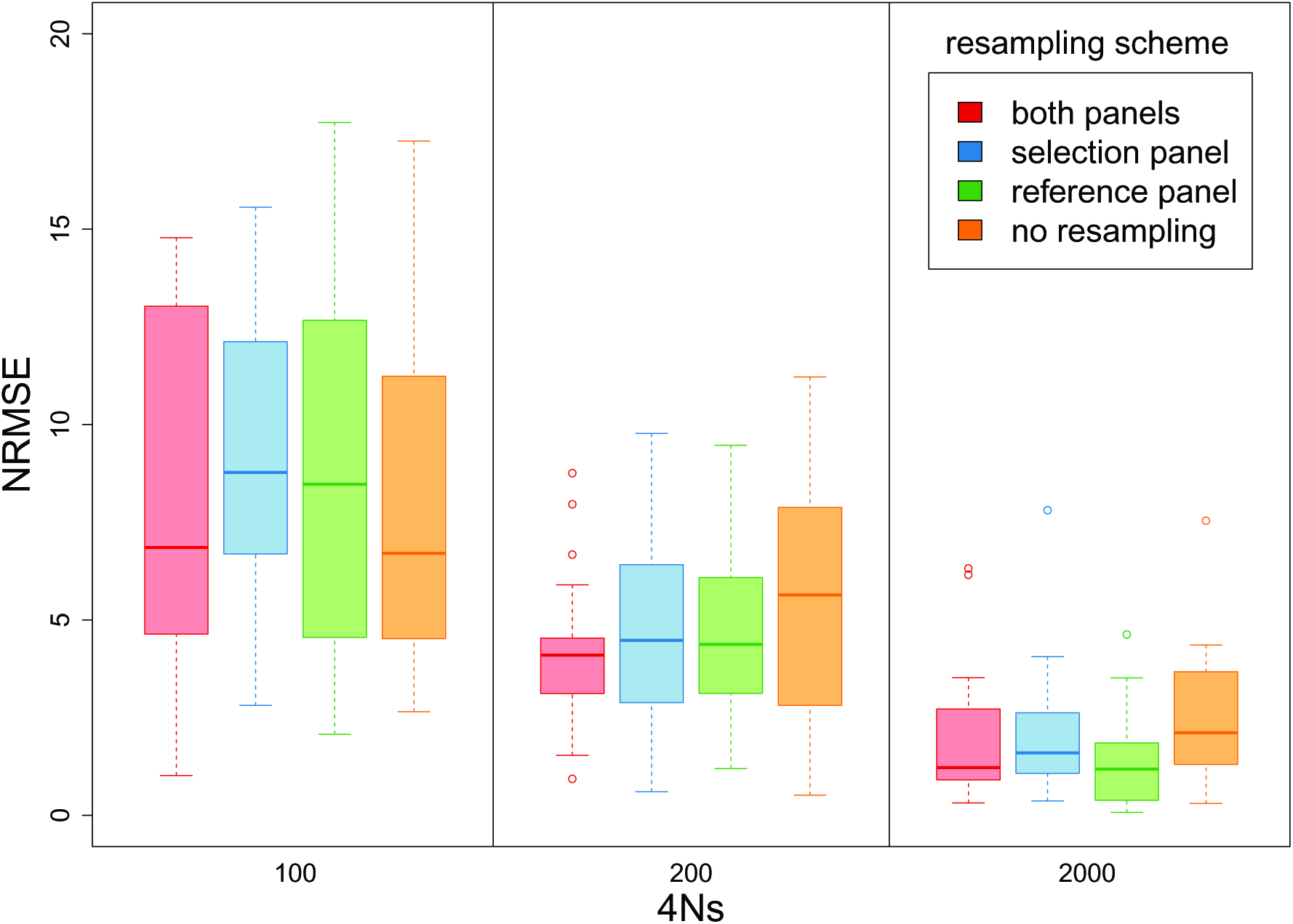
Effects of resampling subsets of complete data. Estimated accuracy and among independent MCMC runs for different resampling schemes. Frequency trajectories were simulated to an end frequency of 0.1. Under each 2Ns value and resampling scheme indicated in the legend, 20 data sets were simulated and inference was performed on the 5 replicate MCMCs. In each simulation, the full dataset includes sample sizes of 100 for the selected and reference panels. Inference for each replicate was then performed on 50 selected haplotypes and 20 reference haplotypes according to the sampling scheme in the legend. Normalized RMSE values are calculated using the estimates and true TMRCA value, while the standard deviations are calculated using the estimates and their mean.

**S6 Fig.**
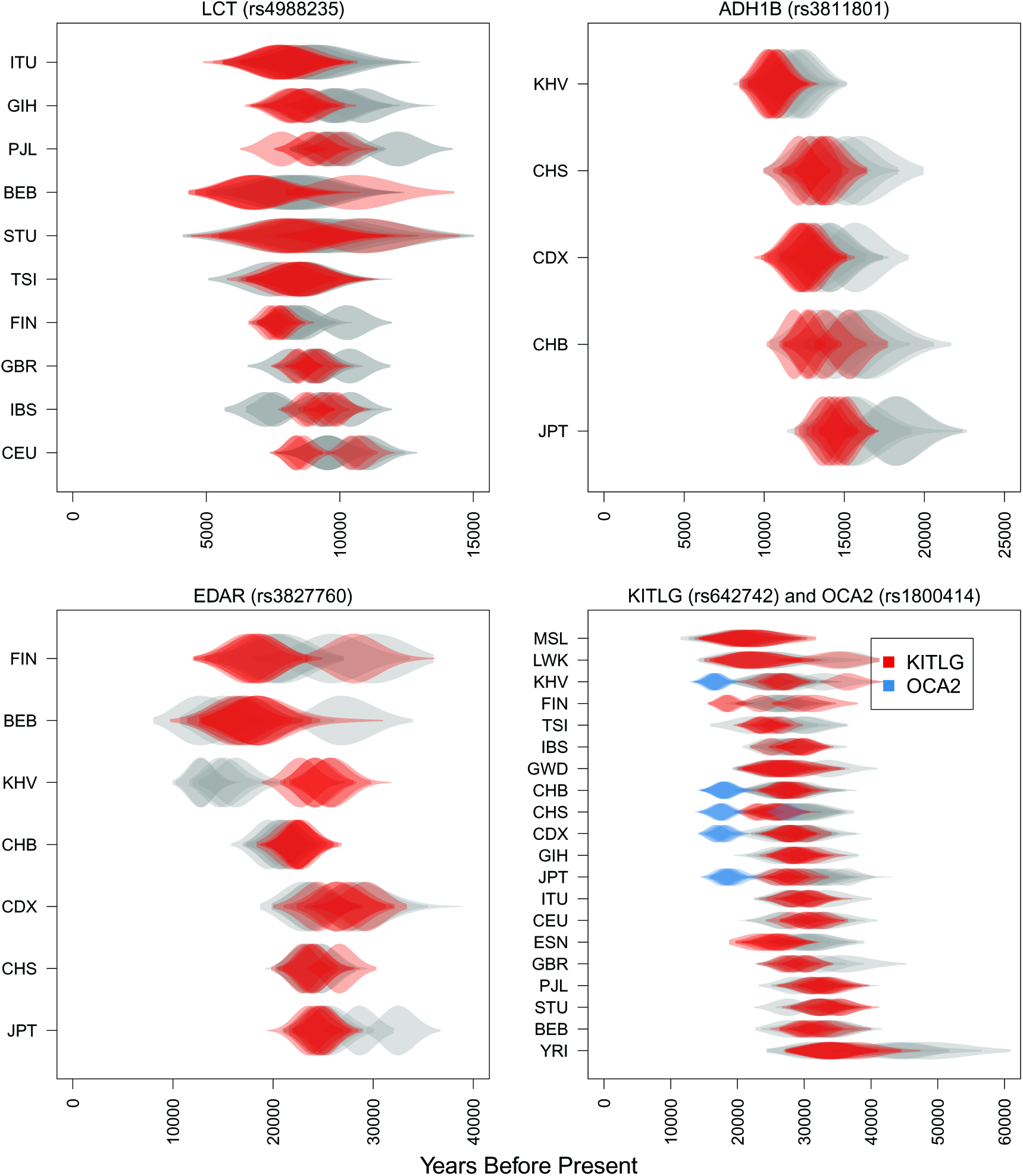
Comparison of fine scale and megabase scale recombination maps. A comparison between estimates made using the fine-scale Decode recombination map (grey) and a uniform recombination rate (red and blue). The uniform recombination rate used for each gene is the mean rate for the 1Mb region around each variant indicated by the rs number. Five replicate MCMCs were performed for each variant and population by resampling the selected and reference panels with replacement. Each MCMC was run for 5000 iterations with a standard deviation of 20 for the t proposal distribution. Replicate MCMCs are plotted with transparency.

## S1 APPENDIX Initializing the ancestral haplotype for the MCMC

To decrease run times for the MCMC, we initialize the starting sequence for the ancestral haplotype using a heuristic algorithm which exploits the decrease in polymorphism near the selected site. Let *A*^0^ denote the initial ancestral haplotype to be estimated, and let the indicator variable *I_ij_* denote whether chromosome *i* is part of the ancestral haplotype at site *j*:

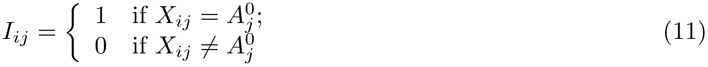

The algorithm proceeds as follows:

1. At *j* = 1 all chromosomes with the beneficial allele are specified to be on the ancestral haplotype at the selected site, i.e. 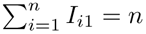 and 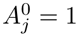.
2. Moving to the next adjacent SNP, we calculate the allele frequency, *F_j_*, among chromosomes on the ancestral haplotype at the previous site:

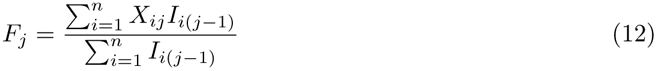
3. The major allele among advantageous allele carriers is assumed to be the putative ancestral allele and minor alleles are assumed to be the result of a putative recombination event off of the ancestral haplotype in the previous SNP interval. For *j* > 0,

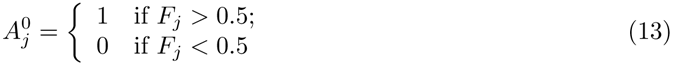

Because we expect there to be some rare or singleton variants on the ancestral haplotype, singletons are removed before step 1 in an effort to improve estimates of the ancestral haplotype at more distant sites. In addition, major and minor alleles can't be identified at sites with alleles at 0.5 frequency and are also removed initially. Steps 2 and 3 are computed iteratively until reaching the end of the locus (*j* = *L*) on both sides flanking the selected site. The sites that were removed (*F^j^* = 0.5 and singletons) are then added back in and take values of *I*_*ij*_ from *I*_*ij*+1_. *A*^0^*_j_* for the added sites are computed using equations 12 and 13. At sites for which 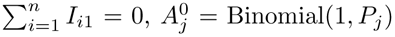, where 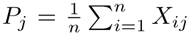. After getting the initial estimate 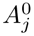, the MCMC is run and evaluated for convergence by visual inspection of trace plots.

**S1 Table 1.** TMRCA estimates from 1000 Genomes Project data. TMRCA estimates from the 1000 Genomes Project panel using the Mb and fine scale recombination rate. These results represent the distributions with the highest posterior probability among the 5 replicates shown with transparency in Figures 6 and 7. All estimates are scaled to a generation time of 29 years.

|  |  | Mb scale recombination map |  | fine scale recombination map |  |
| --- | --- | --- | --- | --- | --- |
| gene | population | $t$ (years) | 95% credible interval | $t$ (years) | 95% credible interval |
| KITLG | FIN | 18733.11 | 16675.03 - 20816.16 | 26343.99 | 21185.68 - 31439.77 |
| KITLG | MSL | 22339.79 | 15723.06 - 28949.85 | 20244 | 15042.13 - 26063.71 |
| KITLG | LWK | 22783.5 | 17921.93 - 28011.88 | 24200.86 | 16839.43 30730.11 |
| KITLG | KHV | 24544.92 | 21643.47 - 27192.74 | 26697.31 | 22249.2 31526.1 |
| KITLG | ESN | 26254.27 | 22854.93 - 29657.6 | 31791.02 | 26440.75 36543.8 |
| KITLG | TSI | 26427.6 | 24109.65 - 28905.17 | 22776.53 | 18379.65 27588.96 |
| KITLG | CHS | 26535.39 | 23456.37 - 29651.06 | 28396.03 | 23284.08 33294.5 |
| KITLG | CHB | 26772.95 | 24297.24 - 30141.32 | 28968.9 | 24451.06 34718.66 |
| KITLG | GBR | 26785.87 | 23841.96 - 29252.6 | 36132.87 | 31410.16 41697.2 |
| KITLG | GWD | 27669.05 | 21900.96 - 33664.19 | 25833.85 | 20134.3 32433.62 |
| KITLG | ITU | 28093.72 | 24607.36 - 31040.55 | 28090.16 | 24212.08 32250.46 |
| KITLG | CDX | 28362.37 | 25128.88 - 31245.57 | 29010.65 | 24457.71 33869.7 |
| KITLG | JPT | 28636.9 | 26351.84 - 31139.23 | 31634.52 | 27551.51 36471.26 |
| KITLG | GIH | 29029.82 | 25862.18 - 32439.4 | 28935.4 | 24599.77 33752.46 |
| KITLG | IBS | 29730.96 | 26169.86 - 32812.62 | 25373.45 | 21393.79 30031.51 |
| KITLG | CEU | 31287.49 | 27866.12 - 34512.88 | 34009.13 | 29818.32 38072.96 |
| KITLG | STU | 32021.3 2 | 8243.9 - 36318.9 | 27693.36 | 23968.81 32516.82 |
| KITLG | BEB | 32030.37 | 29000.59 - 34975.94 | 34375.32 | 29254.91 39578.63 |
| KITLG | PJL | 33719.74 | 30137.36 - 37310.9 | 31384.02 | 26814.24 36005.58 |
| KITLG | YRI | 33947.53 | 28861.11 - 39098.89 | 44437.11 | 36047.45 54074.11 |
| EDAR | FIN | 17386.19 | 13887.25 - 20794.2 | 20176.21 | 15053.08 25838.39 |
| EDAR | BEB | 18370.17 | 14325.16 - 22871.72 | 18418.06 | 13680.78 25409.82 |
| EDAR | CHB | 22192.42 | 19682.73 - 25735.6 | 19262.27 | 16921.98 21521.19 |
| EDAR | JPT | 23508.87 | 21595.24 - 25644.81 | 25730.04 | 23096.63 28826.86 |
| EDAR | CHS | 24058.94 | 22005.79 - 26678.85 | 24813.16 | 22493.15 27204.94 |
| EDAR | CDX | 24360.34 | 21572.05 - 27044.11 | 24346.84 | 21214.9 28019.59 |
| EDAR | KHV | 25683.33 | 23169.98 - 28379.79 | 12686.77 | 11001.3 14645.05 |
| OCA2 | KHV | 16370.39 | 14439.12 - 18102.08 | 26904.93 | 22093.63 32402.26 |
| OCA2 | CHS | 17316.96 | 14913.26 - 19799.16 | 26377.66 | 21217.62 31921.16 |
| OCA2 | CHB | 17838.58 | 15336.98 - 20174.82 | 25159.82 | 20764 29688.86 |
| OCA2 | CDX | 18083.2 1 | 6231.36 - 20253.74 | 28644.18 | 24241.91 33819.6 |
| OCA2 | JPT | 18598.62 | 16110.22 - 20785.6 | 31582.69 | 27875.01 35522.35 |
| ADH1B | KHV | 10841.65 | 9720.032 - 12147.5 | 11186.97 | 9503.454 12862.22 |
| ADH1B | CHS | 12101.84 | 10668.909 - 13479.33 | 15352.24 | 12969.029 17974.85 |
| ADH1B | CDX | 12176.61 | 10678.377 - 13699.32 | 13568.88 | 11183.01 15941.4 |
| ADH1B | JPT | 13996.17 | 12670.869 - 15278.67 | 18317.67 | 15995.495 20911.5 |
| ADH1B | CHB | 15377.36 | 13763.712 - 17281.5 | 13526.93 | 11280.86 16210.89 |
| LCT | BEB | 6869.385 | 5143.203 - 8808.557 | 7971.853 | 5893.793 10443.94 |
| LCT | FIN | 7545.399 | 6982.857 - 8112.515 | 10332.821 | 9349.834 11427.629 |
| LCT | ITU | 7795.401 | 6199.996 - 9419.64 | 8972.475 | 7043.311 11015.566 |
| LCT | TSI | 7936.011 | 6616.676 - 9435.192 | 8630.238 | 7084.033 10230.152 |
| LCT | STU | 8197.625 | 6167.62 - 10338.243 | 7671.266 | 5205.261 10364.956 |
| LCT | GBR | 8412.48 | 7754.023 - 9084.704 | 8185.932 | 7111.164 9226.64 |
| LCT | CEU | 8662.519 | 8064.022 - 9340.064 | 10701.2 | 9579.387 11839.975 |
| LCT | GIH | 8732.25 | 7724.106 - 9921.599 | 9926.97 | 8596.736 11379.234 |
| LCT | IBS | 9341.408 | 8687.717 - 9988.713 | 7593.055 | 6602.516 8681.566 |
| LCT | PJL | 9514.453 | 8596.386 - 10382.874 | 9500.563 | 8511.207 10618.241 |

**S1 Table 2.**
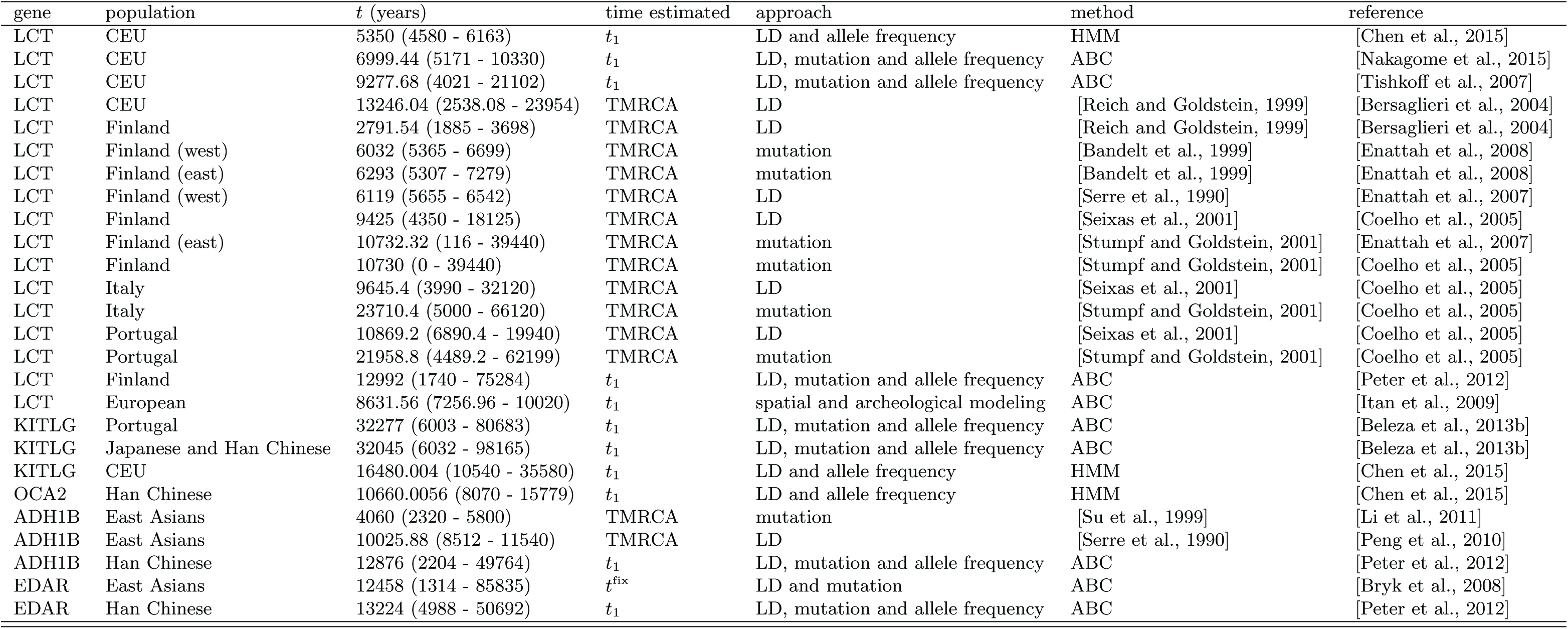
Previous allele age point estimates and 95% confidence intervals for the loci considered in this study. All estimates are scaled to a generation time of 29 years. For the times estimated in each case, *t*_1_ refers to the time of mutation and *t*^fix^ refers to the time since fixation [Przeworski, 2003].

